# Intracellular carbon storage by microorganisms is an overlooked pathway of biomass growth

**DOI:** 10.1101/2022.06.28.497677

**Authors:** Kyle Mason-Jones, Andreas Breidenbach, Jens Dyckmans, Callum C. Banfield, Michaela A. Dippold

## Abstract

The concept of microbial biomass growth is central to microbial carbon (C) cycling and ecosystem nutrient turnover. Growth is usually assumed to occur by cellular replication, despite microorganisms’ capacity to increase biomass by synthesizing storage compounds. Here we examined whether C storage in triacylglycerides (TAGs) and polyhydroxybutyrate (PHB) contribute significantly to microbial biomass growth, under contrasting conditions of C availability and complementary nutrient supply. Together these compounds accounted for 19.1 ± 1.7% to 46.4 ± 8.0% of extractable soil microbial biomass, and revealed up to 279 ± 72% more biomass growth than observed by a DNA-based method alone. Even under C limitation, storage represented an additional 16 – 96% incorporation of added C into microbial biomass. These findings encourage greater recognition of storage synthesis and degradation as key pathways of biomass change and as mechanisms underlying resistance and resilience of microbial communities.

## 1 Introduction

Microbial assimilation of organic resources is a central process in most ecosystems. Soil heterotrophs perform key steps in terrestrial carbon (C) and nutrient cycles, yet how microorganisms use the available organic resources and regulate their allocation to competing metabolic demands remains a subject of research and debate^1–3^. Microbial assimilation of organic C is often conceptualized as “biomass growth”, which is typically envisioned as an increase in microbial abundance, i.e. replicative growth. However, many microorganisms are capable of storage, defined as the accumulation of chemical resources in particular forms or compartments to secure their availability for future use. Various microbial storage compounds are known, including polyhydroxybutyrate (PHB) and triacylglycerides (TAGs)^4,5^. PHB storage is only known among bacteria, while TAGs are used by both bacteria and fungi^6^. Accumulation of storage compounds corresponds to an increase in microbial biomass without replication, and therefore represents an alternative pathway for growth that is not usually considered in the C cycle.

Conventional methods for measuring soil microbial biomass either require extraction into aqueous solution after chloroform fumigation^7^, thereby excluding hydrophobic storage in PHB and TAGs, or measure biomass proxies such as cell membrane lipids or substrate-induced respiration that are not proportional to storage^8,9^. This potential shortcoming is shared with more recent DNA-based measures of microbial growth^10,11^. Biosynthesis of PHB has been demonstrated by compound-specific measurement in soil^12^ and TAGs in marine and soil systems show responsiveness to resource supply consistent with a C-storage function^13,14^. Since “biomass growth” is a cornerstone concept at scales from local ecological stoichiometry to microbially-explicit Earth system models^15,16^, there is a need to assess how severely the omission of storage may bias our understanding of carbon assimilation and utilisation.

Interpretation of storage patterns is facilitated by distinguishing two storage modes, which represent the end-members on a gradient of storage strategies^6,17^. Surplus storage is the accumulation of resources that are available in excess of immediate needs, at little to no opportunity cost, while reserve storage accumulates limited resources at the cost of other metabolic functions. Surplus storage of C would be predicted under conditions of C oversupply, when replicative growth is constrained by other factors such as nutrient limitation. Reserve storage, on the other hand, predicts that storage may also occur under C-limited conditions. Evidence assembled from pure culture studies confirms the operation of both storage modes among microorganisms^6,18–20^. Here we experimentally investigate the importance of microbial storage in soil, and show how storage responses to resource supply and stoichiometry can advance our understanding of resource allocation and microbial biomass growth. We hypothesized as follows:

1. Microbial storage compounds account for a substantial proportion of soil microbial biomass under C-replete, nutrient-limited conditions.
2. Due to low opportunity costs, surplus storage is likely to be quantitatively more significant at the community scale. Therefore, complementary nutrients will suppress storage compound accumulation in favour of replicative growth.
3. Microbial biomass growth is substantially underestimated by neglecting intracellular storage.

Soil microcosms were incubated under controlled conditions, with C availability manipulated through additions of isotopically labelled (^13^C and ^14^C) glucose, which is a common component of plant root exudates and the most abundant product of plant-derived organic matter decomposition^21^. Nutrient supply (N, P, K and S) was manipulated by adding inorganic fertilizers common in agriculture. A fully crossed design included three levels of C addition (zero-C, low-C and high-C; 0, 90 and 400 μg C/g soil) and two levels of nutrient supply (no-nutrient and nutrient-supplemented) with nutrients supplemented at a level predicted to enable full C assimilation under the high-C treatment. CO_2_ efflux was monitored at regular intervals and microcosms were harvested after 24 and 96 hours to determine microbial biomass (by chloroform fumigation-extraction), dissolved organic carbon (DOC), dissolved nitrogen (DN) and the storage compounds PHB and TAGs. In parallel, a set of smaller microcosms (0.5 g soil) was incubated under otherwise identical conditions to measure microbial growth as the incorporation of ^18^O from H_2_ ^18^O into DNA^11^. This method measures the turnover of the microbial population and therefore captures replicative growth better than tracing of specific C substrates^3^. Together these form the first integrated observations of heterotrophic microbial biomass, growth and storage in a natural microbiome, revealing the importance of storage as a resource-use strategy in response to environmental resource supply and element stoichiometry.

## 2 Results and discussion

### 2.1 Microbial nutrient status and CO_2_ efflux

Patterns of soil respiration were in general agreement with past studies of low-molecular weight organic substance utilization in soil^22–24^, and provide interpretive insight into the resource constraints during storage compound synthesis and degradation.

Glucose addition stimulated large increases in CO_2_ efflux (Figure 1), primarily derived from glucose mineralization (Figure 1, inset). Nutrient supplementation barely affected CO_2_ efflux rates from the zero-or low-C additions and for none of these treatments was N availability (measured as dissolved nitrogen) significantly reduced relative to the control at 24 h (Supplementary Figure S1). Thus, C limitation dominated in the zero- and low-C treatments throughout the experimental period, irrespective of nutrient additions.

**Figure 1:**
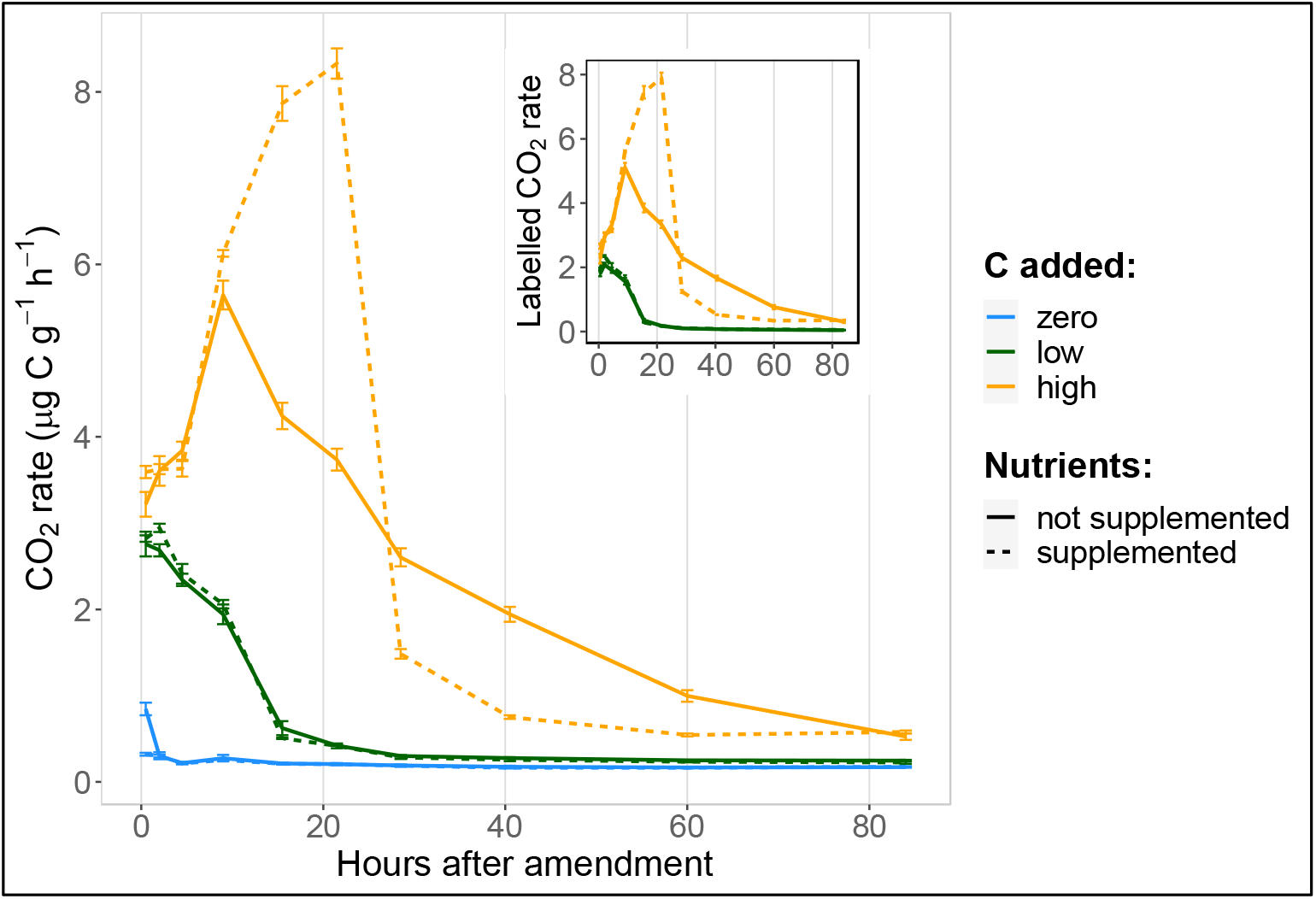
Time-series of the CO_2_ efflux from soil microcosms following addition of a readily degradable ^13^C-labelled carbon source (glucose at 0, 90 and 400 μg C g^-1^ soil) with or without mineral nutrient supply (N, P, K, S). Inset shows only CO_2_ derived from the added glucose in μg C g^-1^ h^-1^. Each point plots the average rate of CO_2_ efflux at the mid-point of the sampling interval, with error bars showing standard deviation (n = 4).

With high-C addition, CO_2_ efflux rates under the two nutrient levels diverged strongly after 12 hours, with the no-nutrient treatment declining steadily from 12 h until the end of the experiment. This early decline in mineralization was consistent with the onset of nutrient limitation, after microbial growth on the added glucose had depleted easily available soil nutrients (Supplementary Figure S2). This depletion was reflected in suppressed dissolved nitrogen after 24 h, with only 35.8 – 62.5% of the zero-C, no-nutrient control (family-wise 95% confidence interval; Supplementary Figure S1) and a further decline to 96 h. Nutrient limitation was accompanied by an accumulation of highly-labelled DOC at 24 h in the soil solution, reflecting unused glucose or soluble by-products in an amount 19.6 ± 2.1% of the original C addition (mean ± standard deviation; Supplementary Figure S3). High C addition without supplementary nutrients resulted in rapid mineralization at first, but nutrient limitation set in within 12 hours and continued for the remaining experimental period.

Nutrient addition had a strong effect in combination with high-C supply: it accelerated glucose mineralization until 24 hours after addition (Figure 1), after which CO_2_ efflux dropped precipitously to below that of the high-C, no-nutrient treatment. Dissolved N decreased only moderately over 24 h (56.2 – 97.9% of control, despite the added nutrients). DOC at 24 h was far lower than in the absence of added nutrients, with no further change to 96 h, despite higher N availability (Cohen’s d >> 1, family-wise p < 0.001 for both), indicating that the microbial community had depleted the added C and re-entered C-limited conditions. Therefore, high C addition with supplementary nutrients maintained rapid C mineralization through the first 24 hours, but glucose depletion then reasserted C-limitation for the rest of the experimental period.

### 2.2 Presence and synthesis of microbial storage compounds

PHB and TAGs were both found in the control soil (zero-C, no nutrients after 24 h; Figure 2, A&C), together representing a pool 24.7 ± 2.5% (mean ± standard deviation) as large as the extractable microbial biomass (Figure S2). This ratio ranged from 19.1 ± 1.7% to 46.4 ± 8.0% over all treatments, indicating that storage is a significant pool of biomass not only under C-replete conditions, as hypothesized, but even when C availability is limited. Storage equivalent to a substantial proportion of biomass offers a resource for regrowth following disturbance, indicating a potential role of storage in supporting resilience of this soil microbial community. Furthermore, most common measures of soil microbial biomass rely on proxies such as soluble carbon after chloroform fumigation, which do not capture this biomass component. This suggest that microbial biomass C may be widely underestimated in soil.

**Figure 2:**
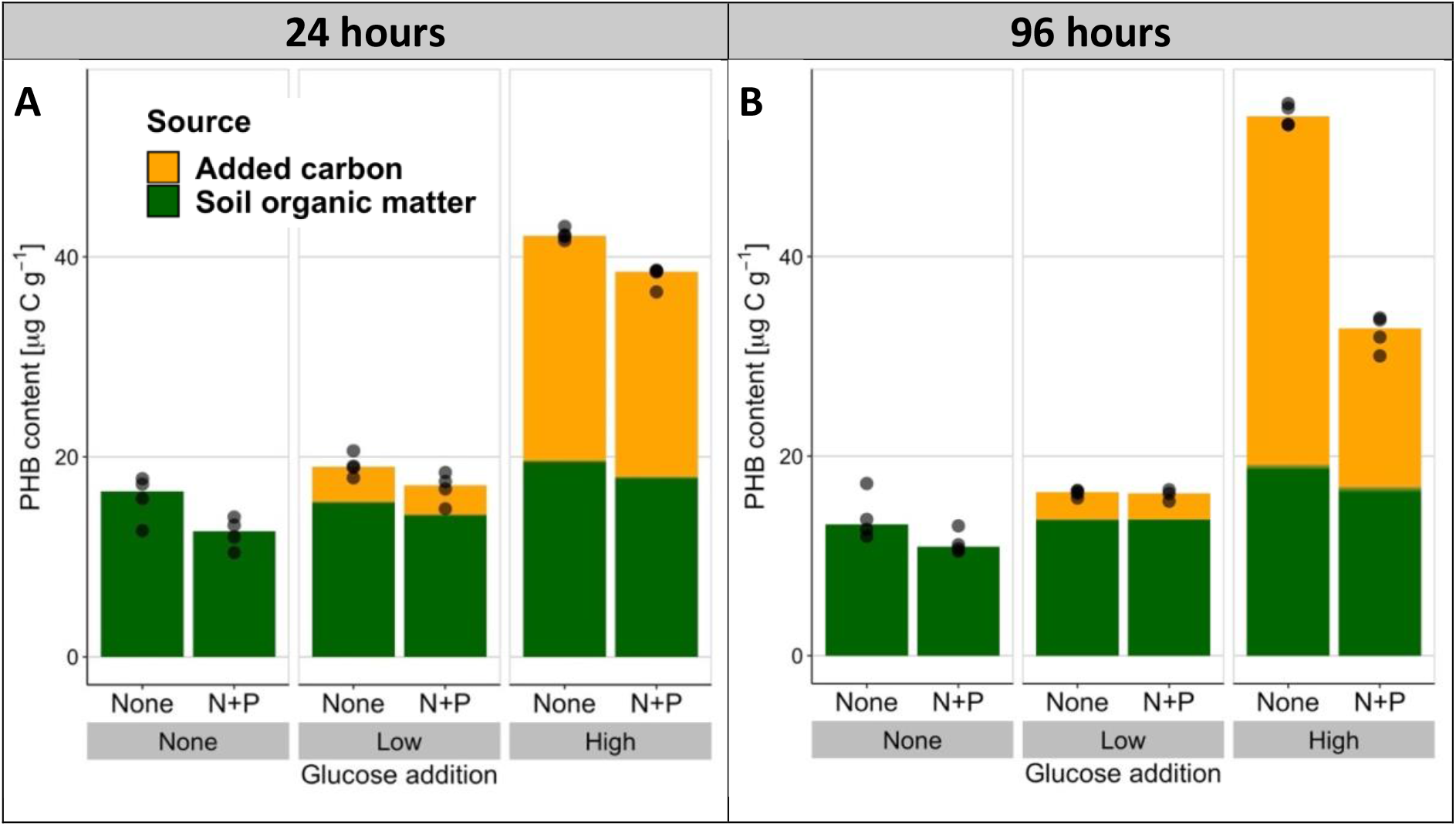
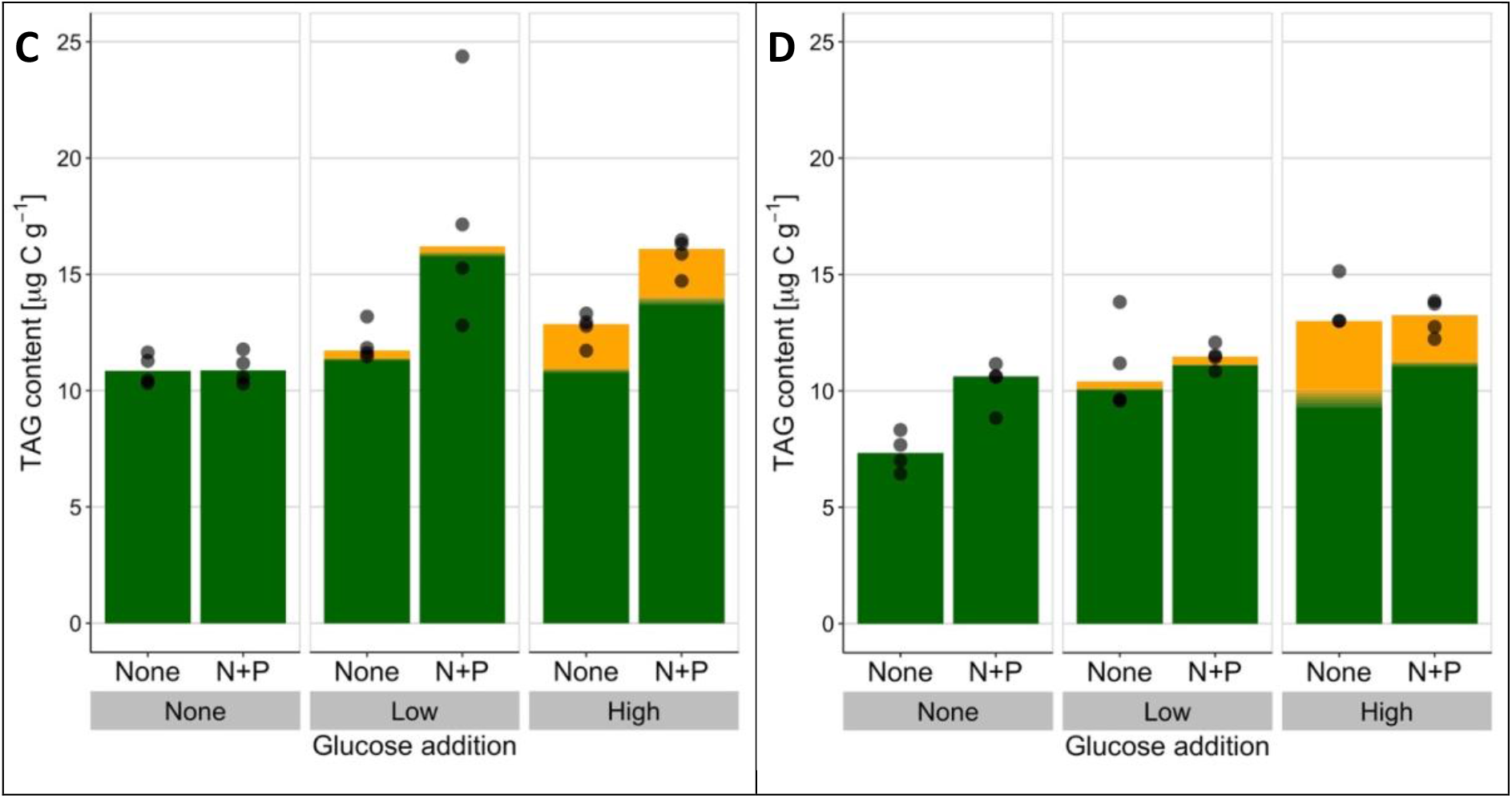
Storage compounds PHB (above, A, B) and TAGs (below, C, D) in soil following addition of a readily degradable, ^13^C-labelled carbon source (glucose at 0, 90 and 400 μg C g^-1^ soil) with or without mineral nutrient supply (N, P, K, S). Soil was sampled after 24 h (left) and 96 h (right). The source of the stored C is shown in contrasting colours as determined by isotopic composition, with uncertainty in relative composition shown as shading of the colours around the mean of composition (shading corresponds to +/-standard deviation, n = 4, except for 1 treatment in each of TAGs and PHB where n = 3).

The two storage compounds were both responsive to the supply of C and complementary nutrients, but with very different behaviours. At both timepoints, the low input of C stimulated only a moderate increase in total PHB, irrespective of nutrient supply. In contrast, high C input stimulated a large increase in PHB, particularly when not supplemented with nutrients (a 308% increase over control at 96 h, with Hodges-Lehmann median difference of 36.0 – 42.9 μg C g^-1^). Nutrient supply significantly suppressed PHB storage, even in the absence of added C (nutrient main effect, robust ANOVA of medians 24 h: F_(1,∞)_ = 35, p < 0.001; 96 h: F_(1,∞)_ = 275, p <0.001). Assimilation of glucose C into new PHB continued between 24 and 96 h under the nutrient-limited conditions of the high-C, no-nutrient treatment (Hodges-Lehmann median difference of 10.2 – 13.3 μg C g^-1^, 95% confidence interval), while in contrast the increasing C limitation of the high-C, nutrient-supplemented treatment after 24 h induced degradation of PHB during this late incubation period (median reduction of 2.7 – 8.6 μg C g^-1^). The PHB storage pool therefore responded dynamically to shifts in resource stoichiometry on a timescale of hours to days, with changes as expected from a surplus storage strategy. Dynamic build-up of storage under conditions of surplus and mobilisation under scarcity mirrors diurnal storage oscillations observed in the ocean^14^ as well as patterns described in pure culture^25^. This study provides the first confirmation of such microbial storage dynamics in a terrestrial ecosystem. At the end of the incubation, stored C was sufficient to completely support basal respiration for 109 – 347 h (depending on the treatment), which could be a crucial resource for withstanding starvation or other stress. Much longer periods would be envisaged if accompanied by strong downregulation of energy use in response to the stress^26^. Thus, the resource buffer provided by storage could help microorganisms in terrestrial ecosystems overcome resource fluctuations and support short-term resistance against environmental disturbance^6^.

Storage of TAGs was enhanced by C input (Figure 2, C&D), but its response to resource stoichiometry differed greatly from PHB. Over 24 h, nutrient supplementation stimulated more TAG accumulation, rather than suppressing it (main nutrient effect F_(1,∞)_ = 10.8, p = 0.001 and nutrient:glucose interactions between zero-C and the two C-supplemented treatments, both p < 0.01), while over 96 h nutrient supply had little effect with C addition and increased TAGs when C was not added (95% confidence interval for median difference 0.5 – 4.7 μg C g^-1^). The TAG response to C and nutrient supply over 96 h resembled changes in extractable microbial biomass (Supplementary Figure S2), which was increased by C supply but not significantly enhanced by nutrients (ANOVA main effect of C supply at 96 h: *F*(2,17) = 8.0, p = 0.003). Therefore, unlike PHB, TAG synthesis was not stimulated by a stoichiometric surplus of available C, suggesting a reserve storage function for this compound. Notably, the relative allocation of glucose C between PHB and TAG remained relatively constant (glucose-derived PHB:TAG ranged 7.0 – 11.5 across all treatments) because the C source used for TAG biosynthesis varied more strongly than total TAG levels in response to C supply. This corroborates a reserve storage function of TAG, with total storage synthesis regulated independently of C supply and drawing on whichever C resources are available, whether glucose-or soil-derived. A reserve storage role for TAG contrasts with an earlier report that fungal TAG accumulation in a forest soil was largely eliminated by complementary nutrient supply^13^, with the difference possibly attributable to the much higher amounts of C provided in that experiment (16 mg glucose-C g^-1^). In our experiment C was traced into both bacterial (16:1ω7) and fungal (18:2ω6,9) TAGs (Supplementary material Figures S4, S5 and S6). The fungal biomarker 18:2ω6,9 was only a minor contributor to TAG incorporation in the current experiment, yet even this fungal TAG was not suppressed by nutrient addition. Our results indicate that both fungi and bacteria employed TAGs as a reserve storage form, with overall levels of TAG storage more closely linked to replicative growth than to resource stoichiometry.

In summary, the response of PHB storage to different C and nutrient conditions was largely consistent with the hypothesized surplus storage mode. In contrast, patterns of TAG storage were better characterized by the reserve storage mode. Since some bacterial taxa can utilize both PHB and TAGs^27,28^, the question arises whether these compounds fulfil different storage functions in individual organisms, or whether the different responses emerge at a community scale, with each compound consistently preferred by a different set of microbial taxa, following divergent storage strategies. In the latter case, storage traits may prove useful contributions to microbial trait-based frameworks as proxies of an organism’s resource allocation strategy.

### 2.3 Microbial storage as a component of biomass growth

The incorporation of C into soil microbial biomass is an essential step in the terrestrial C cycle, and appropriate estimates of these flows are required for C modelling and environmental management. We performed a parallel experiment to measure microbial growth using ^18^O incorporation into DNA^11^. This method is calibrated to units of carbon content based on extractable biomass from the chloroform fumigation-extraction method, and therefore does not capture hydrophobic PHB or TAG storage. We compared the ^18^O-based measure of growth with the net incorporation of isotopically labelled glucose carbon into storage compounds (Figure 3). This provides a comparison of magnitude using a lower bound for storage synthesis by neglecting the biosynthesis of storage from other C sources and any degradation of labelled storage prior to measurement. Furthermore, only two storage forms were measured here, whereas other microbial storage compounds are also known^6^. Storage comprised up to 279 ± 72% more biomass growth than observed by the DNA-based method (mean ± standard deviation for the high-C, no-nutrient treatment at 24 h, Figure 3A). Even under conditions of C limitation, this storage growth represented an additional 16 – 96% incorporation of C into biomass. Intracellular storage evidently plays a quantitatively significant role in microbial assimilation of C under a broad range of stoichiometric conditions, and biomass growth would be substantially underestimated by neglecting storage. As microbial growth is a central variable in microbially-explicit models of the carbon cycle^29^, the substantial scale of storage also encourages a reassessment of model inputs and interpretation of results wherever short-term measurements or dynamic changes are involved.

**Figure 3:**
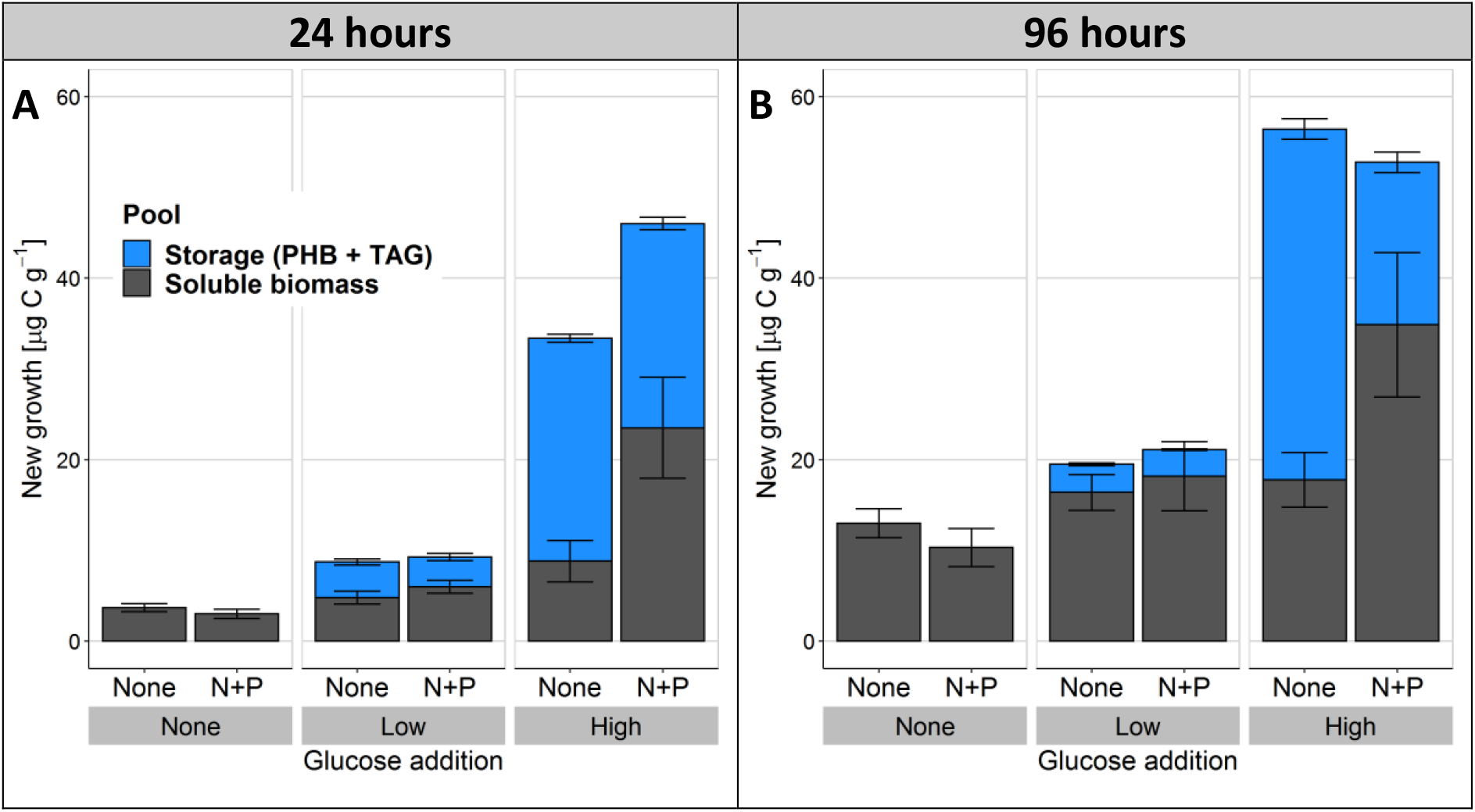
Conservative estimation of new storage biosynthesis in comparison to DNA-based microbial growth reveals storage as a substantial, overlooked component of biomass growth in soil. Here ^13^C-labelled storage compound synthesis (PHB and TAGs) and DNA-based growth (incorporation of ^18^O) were measured in soil over 24 (A) and 96 hours (B) following addition of a readily degradable, ^13^C-labelled carbon source (glucose at 0, 90 and 400 μg C g^-1^ soil) with or without mineral nutrient supply (N, P, K, S). Error bars represent standard deviations in each component of the stacked bar (n = 4).

“Microbial biomass growth” is frequently understood as synonymous with an increase in individuals, in other words, the replicative growth of microbial populations. However, the incorporation of C into storage compounds represents an alternative growth pathway (Figure 4), which differs from replicative growth in crucial ways. Models of microbial growth typically assume that increases in biomass match the elemental stoichiometry of the total biomass (the assumption of stoichiometric homeostasis^30^), and therefore implement overflow respiration of excess C under conditions of C surplus^31^. However, substantial incorporation of C into otherwise nutrient-free PHB and TAG clearly does not follow whole-organism stoichiometry. Growth in storage therefore increases total biomass under stoichiometrically unbalanced conditions. The short experimental timeframe here is representative of environmental resource pulse and depletion processes, such as the arrival of a root tip in a particular soil volume or death and decay of a nearby organism. Storage provides stoichiometric buffering during such transient resource pulses, which is predicted to increase C and N retention over the longer term^32^. By enhancing the efficiency with which microbes incorporate transient resource pulses and supporting metabolic activity through periods of resource scarcity, storage can contribute to resistance and resilience of microbial communities facing environmental disturbances.

**Figure 4:**
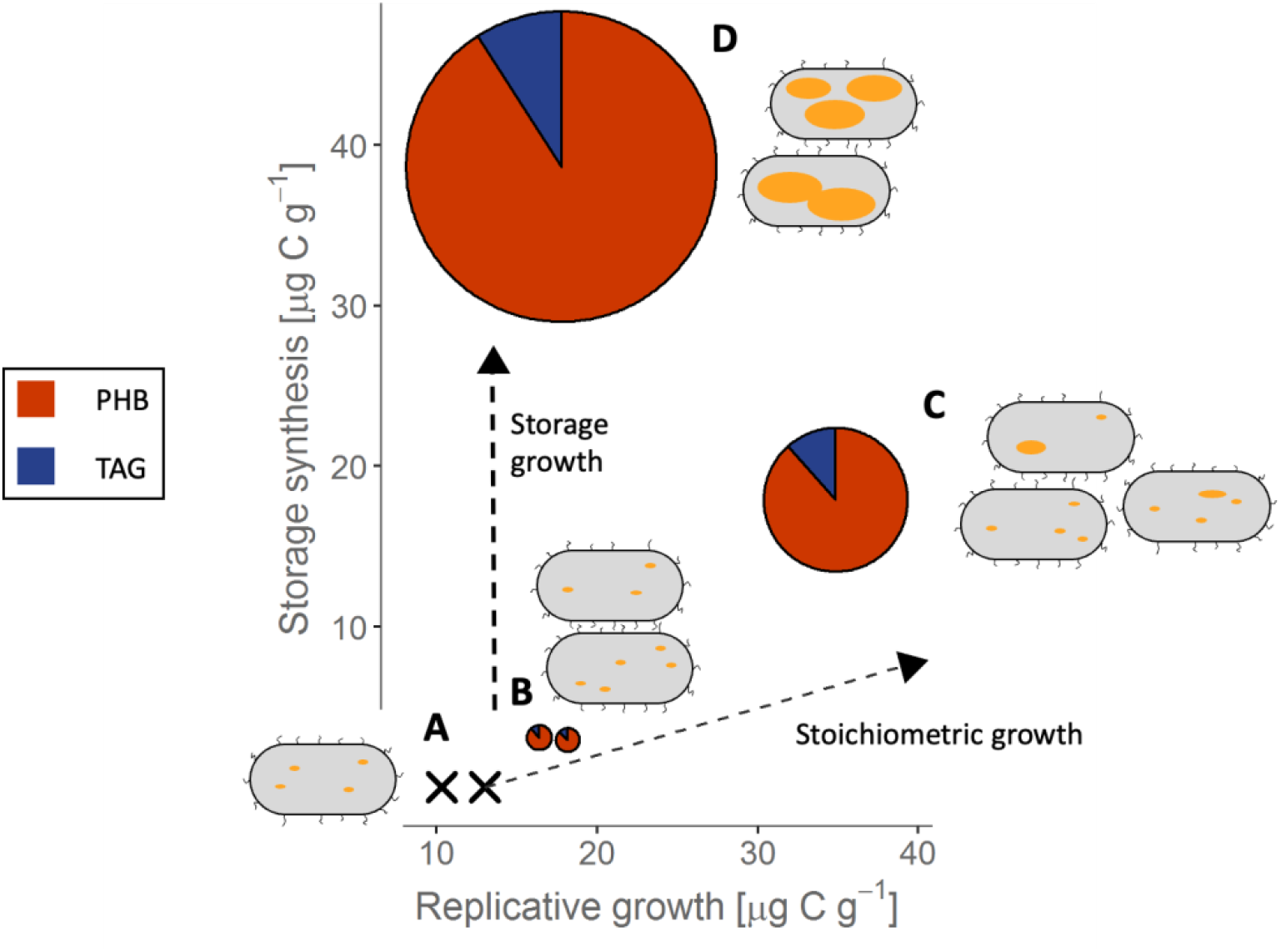
Intracellular storage represents an alternative pathway for growth of microbial biomass, which can be quantitatively substantial but is usually omitted from contemporary discussions. In this conceptual figure the y-coordinates and radii of the pie charts reflect the measured incorporation of added C into storage, with pie chart colours showing the contributions of PHB and TAG (× for zero-C treatments). A microbial population is shown schematically by bacterial cells, with yellow lipid inclusion bodies representing storage. Without C supply, only low levels of replicative growth occur (**A**). Low C additions (with ample nutrients) stimulate replicative growth and limited C incorporation into storage (**B**), with proportions of new storage and non-storage biomass staying close to those predicted by an assumption of constant biomass stoichiometry (dashed line to the right). High C addition with complementary nutrients stimulates both strong replicative growth as well as disproportionately large storage synthesis (**C**) moderately violating the stoichiometric assumption. However, nutrient limitation switches growth strongly towards storage (**D**), incorporating C into biomass without proportionate replicative growth. Replicative growth was measured as ^18^O incorporation into DNA (see also Figure 3).

These findings encourage greater recognition of storage synthesis and degradation as pathways of microbial biomass change in natural communities, in addition to cellular replication. Accounting for microbial storage as a key ecophysiological strategy can enrich our understanding of microbial resource use and its quantitative contribution to global biogeochemical cycles.

## 3 Methods

### 3.1 Experimental design

Topsoil (0 – 25 cm) was collected in November 2019 from the Reinshof experimental farm near Göttingen, Germany (51°29’51.0” N, 9°55’59.0” E) following an oat crop. Five samples along a field transect were mixed to provide a single homogenized soil sample. The soil was a Haplic Luvisol, Ph 5.4 (CaCl_2_), C_org_ 1.4%^33^. Soil was stored at 4°C prior to sieving (2 mm) and then distributed into airtight 100 mL microcosms in laboratory bottles with the equivalent of 25 g dry soil at 48% of water holding capacity (WHC). Four replicates were prepared for each treatment and sampling timepoint. Microcosms were placed in a climate-controlled room at 15°C and preincubated for one week before adding treatment solutions.

Treatment solutions provided glucose as a C source (0, 90 or 400 μg C/g soil) in a fully crossed design with added nutrients (combined (NH_4_)_2_SO_4_ and KH_2_PO_4_) or a no-nutrient control. Glucose treatments contained uniformly isotopically labelled glucose (3 atom% ^13^C and 0.19 kBq ^14^C per microcosm, respectively from Sigma-Aldrich, Munich, Germany and from American Radiolabelled Chemicals, Saint Louis, U.S.A.). The N and P addition was set to be sufficient for the complete utilisation of the C in the high glucose treatment, assuming a C:N:P ratio of 38:5:1 for an agricultural microbial community^34^ and a C-use efficiency of 50%^35^. Addition of the treatment solutions raised the soil moisture to 70% of WHC, after which the microcosms were sealed with air-tight butyl rubber septa and their headspace flushed with CO_2_-free synthetic air. Headspace gas was sampled with a 30 mL gas syringe at regular intervals and collected in evacuated exetainers (Labco, Ceredigion, U.K.) for measurement by gas chromatography – isotope ratio mass spectrometry (GC-Box coupled via a Conflo III interface to a Delta plus XP mass spectrometer, all Thermo Fischer, Bremen, Germany). After gas sampling, the headspace in each microcosm was again flushed with CO_2_-free air.

Microcosms were harvested 24 and 96 hours after application of the treatment solutions. The soil in each microcosm was thoroughly mixed by hand for 30 sec and subsampled for chemical analysis.

### 3.2 Chemical analysis

Extractable microbial biomass was measured by chloroform fumigation extraction ^7,36^. Two 5 g subsamples of moist soil were taken from each microcosm. One was immediately extracted by shaking in 20 mL of 0.05 M K_2_SO_4_ for 1 hour at room temperature, then centrifuged and the supernatant filtered. The other was exposed to a chloroform-saturated atmosphere for 24 hours, after which residual chloroform was removed by repeated evacuation and the fumigated soil was extracted in the same manner as the non-fumigated subsample. Extractable MBC was calculated as the difference in DOC between the fumigated and non-fumigated samples, measured on a Multi N/C 2100S analyser (Analytik Jena, Jena, Germany). Glucose-derived MBC was similarly calculated from the difference in radioactivity (^14^C) of the extracts as measured on a Hidex 300 SL scintillation counter (TDCR efficiency correction, Hidex, Turku, Finland) using Rotiszint Eco Plus scintillation cocktail (Carl Roth, Karlsruhe, Germany). Dissolved nitrogen was determined as total nitrogen in the extracts of the unfumigated soil.

PHB was determined by the method of Mason-Jones et al.^12^, using Soxhlet extraction into chloroform followed by acid-catalysed transesterification in ethanol and GC-MS quantification of the resulting ethyl hydroxybutyrate on a 7890A gas chromatograph (DB1-MS column, 100% dimethyl polysiloxane, 15 m long, inner diameter 0.25 mm, film thickness 0.25 μm), with helium (5.0) as the mobile phase at a flow rate of 1 mL min^-1^, coupled to a 7000A triple quadrupole mass spectrometer (all Agilent, Waldbronn, Germany). Injection volume was 1 μL at an inlet temperature of 270°C and split ratio of 25:1. The GC temperature was: 42°C isothermal for 7 min; ramped to 77°C at 5°C min^-1^; then to 155°C at 15°C min^-1^; held for 15 min; and the ramped to 200°C at 10°C min^-1^. The transfer line temperature was 280°C, with electron ionization at 70 eV. Quantification was based on ions at m/z 43, 60 and 87 for the ethyl 3-hydroxybutyrate analyte, and at m/z 57, 71 and 85 for the undecane internal standard. Identity and purity of peaks was confirmed by scan measurement across the range m/z 40 to 300. The same chromatographic conditions were used for determination of the PHB isotopic composition on a Thermo GC Isolink coupled with a Conflo IV interface to a MAT 253 isotope ratio mass spectrometer (all Thermo Fisher, Bremen, Germany), but with splitless injection. The measured isotopic compositions were corrected for carbon added in derivatization according Glaser and Amelung ^37^.

TAGs were extracted using established protocols for neutral and phospholipid analysis in soil ^38^, according to which lipids were first extracted from frozen soil into a single-phase chloroform-methanol-water solution, purified by solvent extraction, and neutral lipids separated from more polar lipids on a silica solid-phase extraction column. Following removal of the solvent by evaporation, the purified TAGs were hydrolyzed (0.5 M NaOH in MeOH, 10 minutes at 100°C) and methylated (12.5 M BF3 in MeOH, 15 minutes at 85°C), followed by extraction into hexane, drying and redissolution in toluene. The resulting fatty acid methyl esters were quantified by GC-MS on a 7890A gas chromatograph (DB-5 MS column, 5%-phenyl methylpolysiloxane, 30 m coupled to a DB1-MS 15m long, both with an inner diameter 0.25 mm and film thickness 0.25 μm) with an injection volume of 1 μl into the splitless inlet heated to 270°C, and at a constant flow of He (4.6) of 1.2 mL min^-1^, coupled to a 5977B series mass spectrometer (Agilent, Waldbronn, Germany), set to 70 eV electron impact energy, with the GC oven programme as follows: initial temperature 80°C isothermal for 1 min, ramped at 10°C min^1^ to 171°C, ramped at 0.7°C min^1^ to 196°C isothermal for 4 min, ramped at 0.5°C to 206°C, and ramped at 10°C min^1^ to the final temperature of 300°C, isothermal for 10 min for column reconditioning. Isotopic composition was determined in triplicate using a Trace GC 2000 (CE Instruments ThermoQuest Italia, S.p.A), coupled with a Combustion Interface III to a DeltaPlus isotope-ratio mass spectrometer (Thermo Finnigan, Bremen, Germany) using the same dimensions and parameters with splitless injection.

Growth was estimated by ^18^O incorporation into DNA^10,11^. Parallel microcosms were prepared with 0.50 g soil in 2 mL Eppendorf tubes (Eppendorf, Hamburg, Germany) and incubated alongside the larger microcosms. Treatment solutions were prepared at the same concentrations as for the larger microcosms, but enriched with 97 atom% H_2_^18^O so that addition provided a final soil solution of 4.2 atom% ^18^O. Tubes were withdrawn from incubation 24 h and 96 h after addition and immediately frozen at -80°C. DNA was subsequently extracted using MP Bio FastDNA Spin Kit for Soil (MP Biomedicals, Solon, OH, USA), following the manufacturer’s recommendations. DNA concentration in the extract was measured on an Implen MP80 nanophotometer (Implen, Munich, Germany) at 260 nm, with A260/280 and A260/A230 to confirm quality, and 50 μL was pipetted into silver capsules, freeze dried, and measured by TC/EA (Thermo Finnigan, Bremen, Germany) coupled with a Conflo III interface to a Delta V Plus isotope ratio mass spectrometer (all Thermo Finnigan, Bremen, Germany). The total measured O content of the sample, the O content of the DNA (31 % by mass), and the ^18^O natural abundance of unlabelled control samples were used to calculate the background ^18^O from the kit. This background ^18^O was deducted to obtain ^18^O abundance of the DNA, which was applied in a 2-pool mixing model with 70% of O in new DNA derived from water^39^. This provided the fraction of extracted DNA that had been newly synthesized during the incubation period. This fraction was multiplied by extractable microbial biomass to arrive at gross biomass growth in units of μg C g^-1^ soil.

### 3.3 Statistical analysis

Statistical analysis was performed in R^40^. Dissolved nitrogen and DOC data was log-transformed to satisfy assumptions for ANOVA (Shapiro-Wilk’s test of normality and Levene’s test for homogeneity of variance), followed by Tukey’s HSD test for pairwise comparisons of treatment effects. Where relevant, effect sizes were computed as Cohen’s d, using the effsize package^41^. The same analysis was performed on untransformed extractable microbial biomass data.

Levels of labelled storage compounds showed considerable heteroskedasticity that could not be consistently corrected by transformation, particularly due to very high levels of unsaturated fatty acids in one of the 24 h samples. This conceivably reflected a hotspot of fungal activity in the soil. This datapoint was therefore conservatively retained since this would comprise relevant variability in the soil. Analysis of storage compounds proceeded by robust ANOVA of medians for each timepoint separately using the R package WRS2^42^. Pairwise tests of median differences in storage (2-sided) were calculated as 95% confidence intervals using the Hodges-Lehmann estimator (R package DescTools^43^).

Growth estimation by ^18^O incorporation used DNA concentration and its ^18^O enrichment to determine mean gross microbial growth for each treatment in relative terms. The corresponding mean extractable microbial biomass values were applied to convert to absolute units of μg C, using standard rules of error propagation^44^.

Results are presented as mean ± standard deviation unless otherwise noted.

## 4 Acknowledgements

We gratefully acknowledge the laboratory assistance provided by Kali Middleby, Andrew Gall, Lydia Köbele and Karin Schmidt. KMJ thanks the Deutscher Akademischer Austauschdienst (DAAD) for a fellowship supporting the experimental work. This work is associated to the DFG Priority Program 2322 “SoilSystems”, project EcoEnergeticS (DFG DI 2136/17-1). We thank the staff of the core projects of the SPP and the scientific committee for establishing the SPP project.

## 5 Author contributions

KMJ, AB, CCB and MAD jointly initiated and designed the experiment; KMJ, AB and JD conducted the experiment and analyses; KMJ and AB undertook data analysis; KMJ wrote the manuscript with assistance and comments of all co-authors.

## References

1. Sokol, N. W. et al. Life and death in the soil microbiome: how ecological processes influence biogeochemistry. Nat Rev Microbiol (2022) doi:10.1038/s41579-022-00695-z.

2. Dijkstra, P. et al. High carbon use efficiency in soil microbial communities is related to balanced growth, not storage compound synthesis. Soil Biol. Biochem. 89, 35–43 (2015).

3. Geyer, K. M., Dijkstra, P., Sinsabaugh, R. & Frey, S. D. Clarifying the interpretation of carbon use efficiency in soil through methods comparison. Soil Biology and Biochemistry 128, 79–88 (2019).

4. Murphy, D. J. The dynamic roles of intracellular lipid droplets: From archaea to mammals. Protoplasma 249, 541–585 (2012).

5. López, N. I., Pettinari, M. J., Nikel, P. I. & Méndez, B. S. Polyhydroxyalkanoates: Much more than biodegradable plastics. in Advances in Applied Microbiology vol. 93 73–106 (Elsevier, 2015).

6. Mason-Jones, K., Robinson, S. L., Veen, G. F., Manzoni, S. & van der Putten, W. H. Microbial storage and its implications for soil ecology. ISME J (2021) doi:10.1038/s41396-021-01110-w.

7. Vance, E. D., Brookes, P. C. & Jenkinson, D. S. An extraction method for measuring soil microbial biomass C. Soil Biol. Biochem. 19, 703–707 (1987).

8. Anderson, J. P. E. & Domsch, K. H. A physiological method for the quantitative measurement of microbial biomass in soils. Soil Biol. Biochem. 10, 215–221 (1978).

9. Zelles, L., Bai, Q. Y., Rackwitz, R., Chadwick, D. & Beese, F. Determination of phospholipid- and lipopolysaccharide-derived fatty acids as an estimate of microbial biomass and community structures in soils. Biol. Fertil. Soils 19, 115–123 (1995).

10. Blazewicz, S. J. & Schwartz, E. Dynamics of ^18^O incorporation from H_2_ ^18^O into soil microbial DNA. Microb Ecol 61, 911–916 (2011).

11. Spohn, M., Klaus, K., Wanek, W. & Richter, A. Microbial carbon use efficiency and biomass turnover times depending on soil depth – Implications for carbon cycling. Soil Biology and Biochemistry 96, 74–81 (2016).

12. Mason-Jones, K., Banfield, C. C. & Dippold, M. A. Compound-specific ^13^C stable isotope probing confirms synthesis of polyhydroxybutyrate by soil bacteria. Rapid Commun. Mass Spectrom. 33, 795–802 (2019).

13. Bååth, E. The use of neutral lipid fatty acids to indicate the physiological conditions of soil fungi. Microb. Ecol. 45, 373–383 (2003).

14. Becker, K. W. et al. Daily changes in phytoplankton lipidomes reveal mechanisms of energy storage in the open ocean. Nat. Commun. 9, (2018).

15. Manzoni, S. & Porporato, A. Soil carbon and nitrogen mineralization: Theory and models across scales. Soil Biol. Biochem. 41, 1355–1379 (2009).

16. Wieder, W. R. et al. Explicitly representing soil microbial processes in Earth system models: Soil microbes in Earth system models. Global Biogeochem. Cycles 29, 1782–1800 (2015).

17. Chapin, F. S., Schulze, E. & Mooney, H. A. The ecology and economics of storage in plants. Annu. Rev. Ecol. Syst. 21, 423–447 (1990).

18. Matin, A., Veldhuis, C., Stegeman, V. & Veenhuis, M. Selective advantage of a Spirillum sp. in a carbon-limited environment. Accumulation of poly-β-hydroxybutyric acid and its role in starvation. J. Gen. Microbiol. 112, 349–355 (1979).

19. Poblete-Castro, I. et al. The metabolic response of P. putida KT2442 producing high levels of polyhydroxyalkanoate under single- and multiple-nutrient-limited growth: Highlights from a multi-level omics approach. Microb. Cell Fact. 11, 34 (2012).

20. Kourmentza, C. et al. Recent advances and challenges towards sustainable polyhydroxyalkanoate (PHA) production. Bioengineering 4, 55 (2017).

21. Gunina, A. & Kuzyakov, Y. Sugars in soil and sweets for microorganisms: Review of origin, content, composition and fate. Soil Biology and Biochemistry 90, 87–100 (2015).

22. Blagodatskaya, E. V., Blagodatsky, S. A., Anderson, T.-H. & Kuzyakov, Y. Priming effects in Chernozem induced by glucose and N in relation to microbial growth strategies. Applied Soil Ecology 37, 95–105 (2007).

23. Schneckenberger, K., Demin, D., Stahr, K. & Kuzyakov, Y. Microbial utilization and mineralization of [14C]glucose added in six orders of concentration to soil. Soil Biology and Biochemistry 40, 1981–1988 (2008).

24. Creamer, C. A., Jones, D. L., Baldock, J. A. & Farrell, M. Stoichiometric controls upon low molecular weight carbon decomposition. Soil Biology and Biochemistry 79, 50–56 (2014).

25. Sekar, K. et al. Bacterial glycogen provides short-term benefits in changing environments. Appl. Environ. Microbiol. 86, e00049–20 (2020).

26. Dijkstra, P. et al. On maintenance and metabolisms in soil microbial communities. Plant Soil (2022) doi:10.1007/s11104-022-05382-9.

27. Kalscheuer, R., Wältermann, M., Alvarez, H. & Steinbüchel, A. Preparative isolation of lipid inclusions from Rhodococcus opacus and Rhodococcus ruber and identification of granule-associated proteins. Archives of Microbiology 177, 20–28 (2001).

28. Alvarez, H. M. Relationship between β-oxidation pathway and the hydrocarbon-degrading profile in actinomycetes bacteria. Int. Biodeterior. Biodegradation 52, 35–42 (2003).

29. Wieder, W. R. et al. Carbon cycle confidence and uncertainty: Exploring variation among soil biogeochemical models. Global Change Biology 24, 1563–1579 (2018).

30. Mooshammer, M., Wanek, W., Zechmeister-Boltenstern, S. & Richter, A. Stoichiometric imbalances between terrestrial decomposer communities and their resources: Mechanisms and implications of microbial adaptations to their resources. Front. Microbiol. 5, (2014).

31. F Wutzler, T., Zaehle, S., Schrumpf, M., Ahrens, B. & Reichstein, M. Adaptation of microbial resource allocation affects modelled long term soil organic matter and nutrient cycling. Soil Biology and Biochemistry 115, 322–336 (2017).

32. Manzoni, S. et al. Intracellular Storage Reduces Stoichiometric Imbalances in Soil Microbial Biomass – A Theoretical Exploration. Front. Ecol. Evol. 9, 714134 (2021).

33. Ehlers, W., Werner, D. & Mähner, T. Wirkung mechanischer Belastung auf Gefüge und Ertragsleistung einer Löss-Parabraunerde mit zwei Bearbeitungssystemen. Journal of Plant Nutrition and Soil Science 163, 321–333 (2000).

34. Xu, X., Thornton, P. E. & Post, W. M. A global analysis of soil microbial biomass carbon, nitrogen and phosphorus in terrestrial ecosystems: Global soil microbial biomass C, N and P. Global Ecol. Biogeogr. 22, 737–749 (2013).

35. Manzoni, S., Taylor, P., Richter, A., Porporato, A. & Ågren, G. I. Environmental and stoichiometric controls on microbial carbon-use efficiency in soils: Research review. New Phytologist 196, 79–91 (2012).

36. Gunina, A., Dippold, M. A., Glaser, B. & Kuzyakov, Y. Fate of low molecular weight organic substances in an arable soil: From microbial uptake to utilisation and stabilisation. Soil Biology and Biochemistry 77, 304–313 (2014).

37. Glaser, B. & Amelung, W. Determination of ^13^C natural abundance of amino acid enantiomers in soil: methodological considerations and first results. Rapid Communications in Mass Spectrometry 16, 891–898 (2002).

38. Banfield, C. C., Dippold, M. A., Pausch, J., Hoang, D. T. T. & Kuzyakov, Y. Biopore history determines the microbial community composition in subsoil hotspots. Biology and Fertility of Soils 53, 573–588 (2017).

39. Pold, G., Domeignoz-Horta, L. A. & DeAngelis, K. M. Heavy and wet: The consequences of violating assumptions of measuring soil microbial growth efficiency using the 18O water method. Elementa: Science of the Anthropocene 8, 069 (2020).

40. F R Core Team. R: A Language and Environment for Statistical Computing. (R Foundation for Statistical Computing, 2020).

41. Torchiano, M. Effsize - A Package For Efficient Effect Size Computation. (Zenodo, 2016). doi:10.5281/ZENODO.1480624.

42. Mair, P. & Wilcox, R. Robust statistical methods in R using the WRS2 package. Behav Res 52, 464–488 (2020).

43. Signorell, A. & et al. DescTools: Tools for descriptive statistics. (2021).

44. Meyer, S. L. Data Analysis for Scientists and Engineers. (John Wiley and Sons, 1975).

