## Supplementary information for "Intracellular carbon storage by microorganisms is an overlooked pathway of biomass growth"

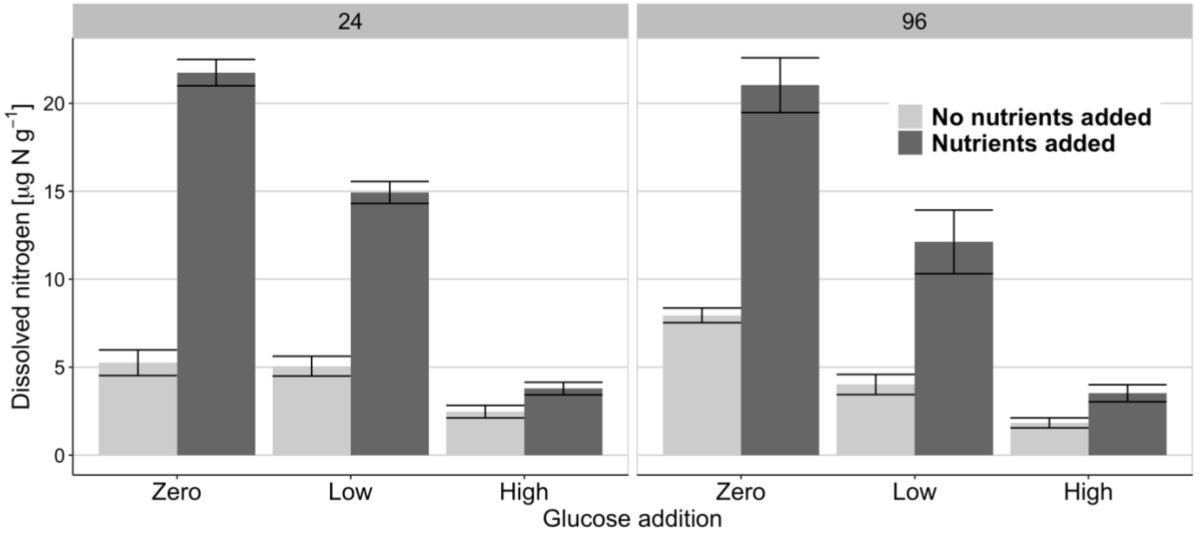


**Figure S1:** Dissolved nitrogen extractable into aqueous 0.05 M $K_2SO_4$

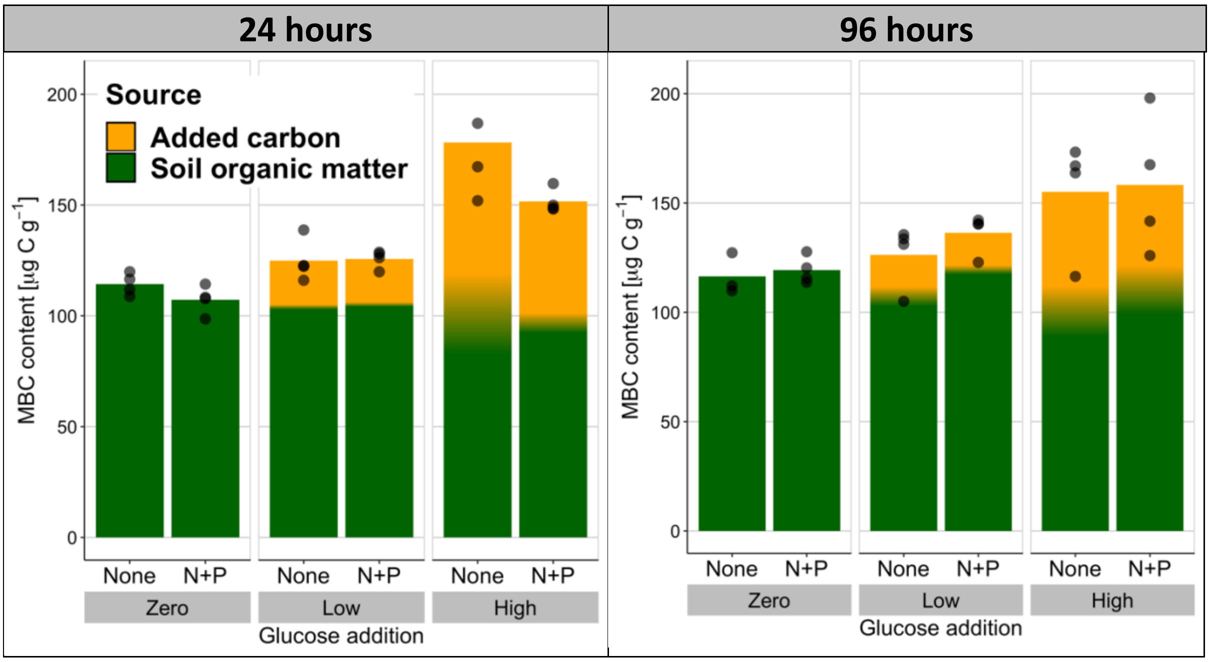


**Figure S2:** Extractable soil microbial biomass determined by chloroform fumigation-extraction. The source of the stored C is shown in contrasting colours as determined by isotopic composition, with shading representing +/- 1 standard deviation (n = 4).

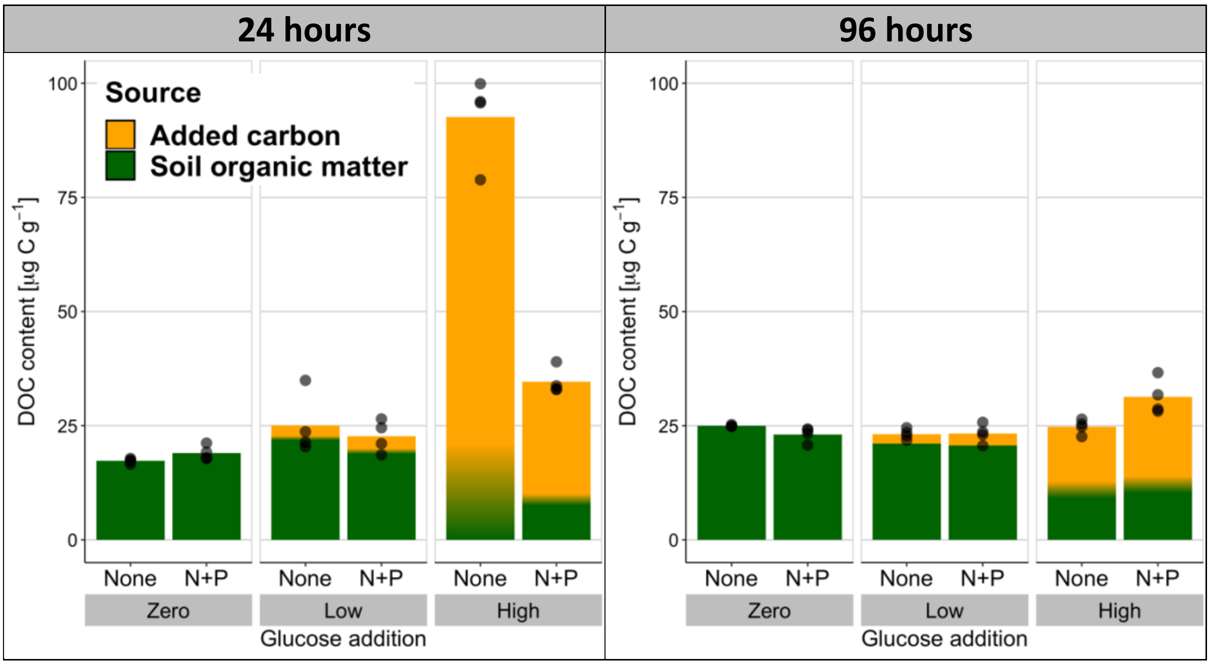


**Figure S3:** Dissolved organic carbon determined by following extraction with 0.5 M $K_2SO_4$. The source of the stored C is shown in contrasting colours as determined by isotopic composition, with shading representing +/- 1 standard deviation (n = 4).

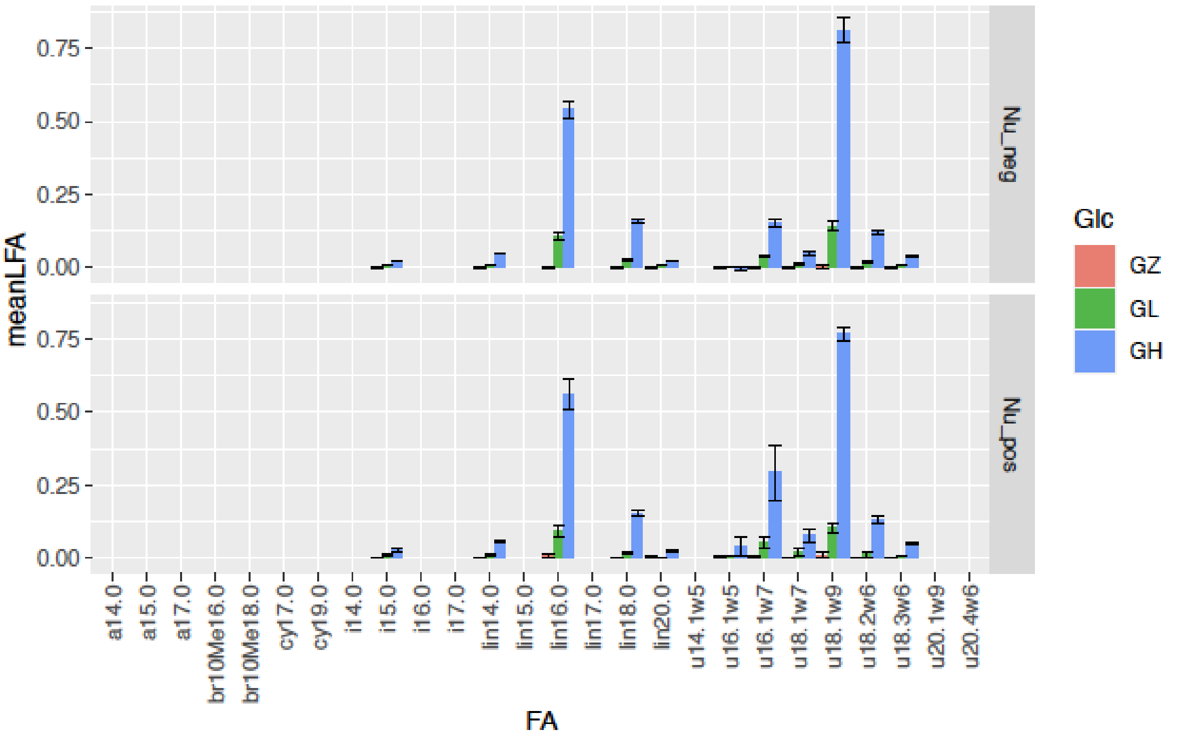


**Figure S4:** Fatty acid profile of glucose-derived TAGs, 24 h after addition of glucose without and with supplementary nutrients (above and below, respectively). Error bars representing +/- 1 standard deviation (n = 4).

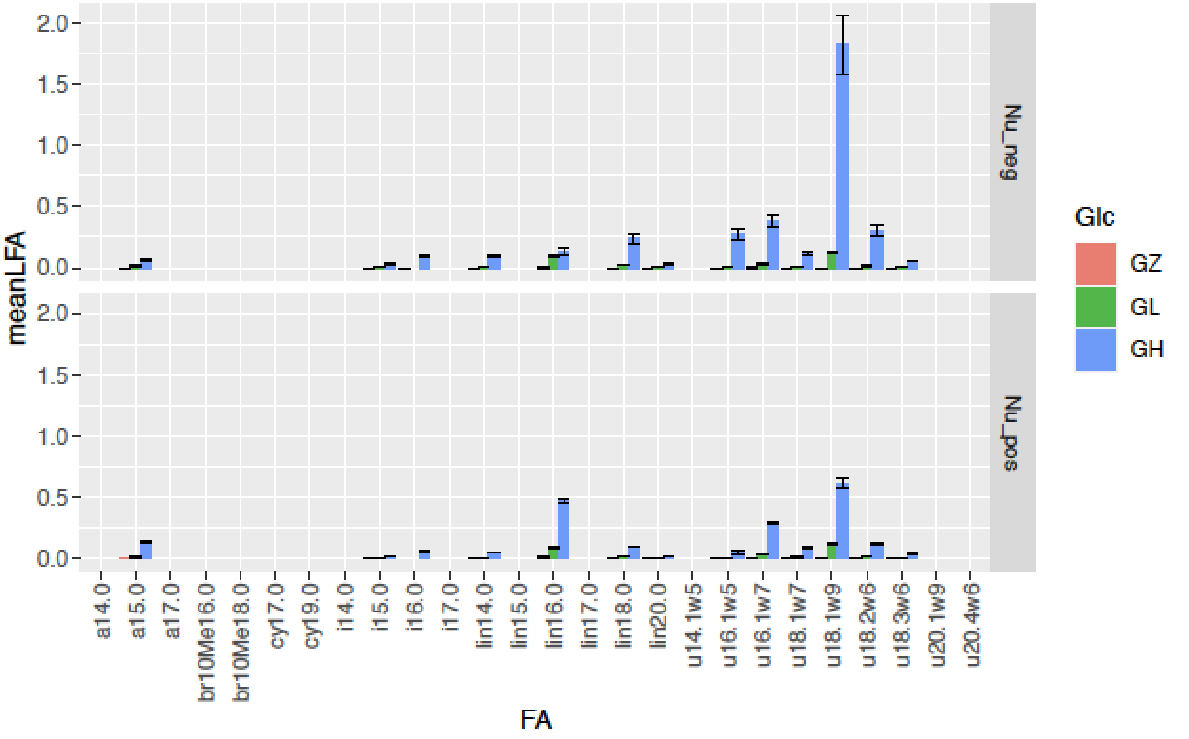


**Figure S5:** Fatty acid profile of glucose-derived TAGs, 96 h after addition of glucose without and with supplementary nutrients (above and below, respectively). Error bars representing +/- 1 standard deviation (n = 4).

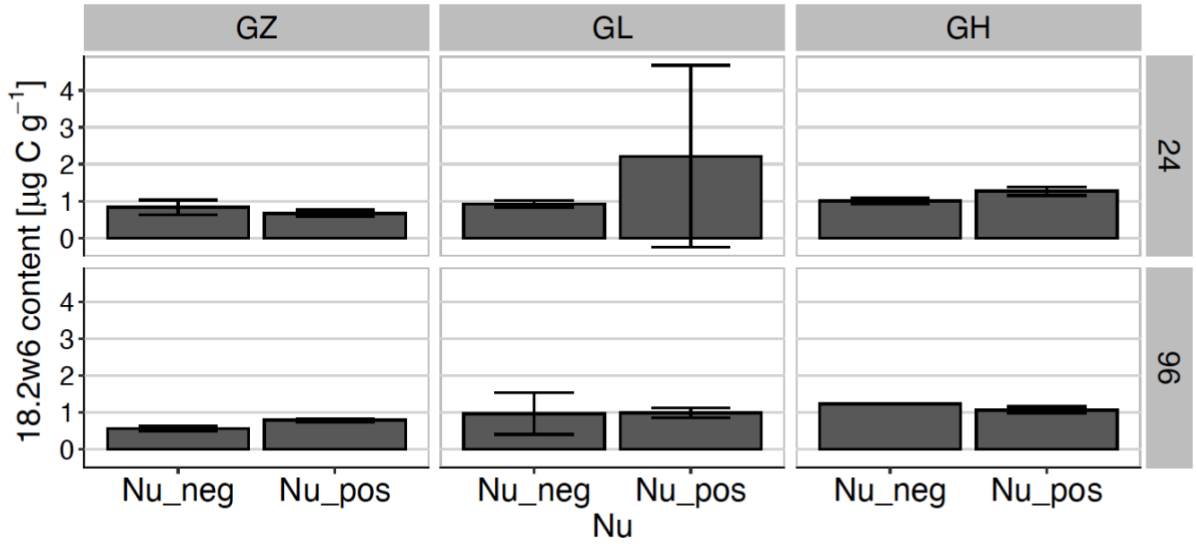


**Figure S6:** Soil content of fungal biomarker TAG 18:2ω6,9. Error bars representing +/- 1 standard deviation (n = 4).
